# Single Cell RNA-Seq Analysis of Regenerative Drug-Treated Human Pancreatic Islets Identifies A Cycling Alpha Cell Population As Key Beta Cell Progenitors

**DOI:** 10.1101/2023.09.07.556578

**Authors:** Esra Karakose, Xuedi Wang, Peng Wang, Saul Carcamo, Deniz Demircioglu, Luca Lambertini, Olivia Wood, Randy Kang, Geming Lu, Donald K. Scott, Adolfo Garcia-Ocaña, Carmen Argmann, Robert Sebra, Dan Hasson, Andrew F. Stewart

## Abstract

Diabetes ultimately results from an inadequate number of functional, insulin-producing human beta cells. Although current attempts to replenish the remaining beta cell pool in people with diabetes are encouraging, scalability and cost limit access for the millions of people with diabetes. The small molecule DYRK1A inhibitor class of beta cell regenerative drugs, either alone or in combination with GLP1 receptor agonists or TGFβ superfamily inhibitors, are capable of inducing beta cell replication *in vitro* and increasing beta cell mass *in vivo*. Despite these advances, the precise mechanisms of action of DYRK1A inhibitors remain incompletely understood. To address the mechanisms more deeply, we performed single cell RNA sequencing on human pancreatic islets treated with a DYRK1A inhibitor, either alone, or in combination with a GLP1 receptor agonist or a TGFβ superfamily inhibitor. We identify a cluster of Cycling Alpha Cells as the cells most responsive to DYRK1A inhibition. Velocity and pseudotime lineage trajectory analyses suggest that Cycling Alpha Cells serve as the primary target cell type for of DYRK1A inhibitors, and may serve as precursor cells that transdifferentiate into functional human beta cells in response to the DYRK1A inhibition. In addition to providing a novel mechanism of action for DYRK1A inhibitors, our findings suggest that efforts to target regenerative drugs to human beta cells may be mis-directed: the proper target may be Cycling Alpha Cells.

## Introduction

Diabetes affects 537 million people globally^1^. While Type 1 diabetes (T1D) and Type 2 diabetes (T2D) differ in their etiology, they share an important common feature; a marked reduction in the number of functional, insulin-producing pancreatic beta cells, suggesting that beta cell replenishment would be of therapeutic value in both diseases. Indeed, strategies to expand the beta cell pool in T1D and T2D are currently in use or in development, including whole pancreas transplantation, pancreatic islet transplantation, and transplant of human stem cell-derived beta cells^2^. Although great progress in these approaches has been made, none is scalable to the millions of people with diabetes, due to high cost, scarcity of donor organ, and their intrinsic interventional nature. These considerations have prompted efforts to develop drugs that may be capable of increasing endogenous beta cell mass. To this end, progress has been made with the discovery of the small molecule drugs that inhibit the, <u>D</u>ual T<u>y</u>rosine-<u>R</u>egulated <u>K</u>inase 1A (DYRK1A), when used alone^3-7^, or in combination with its synergistic partners, GLP1 receptor agonists (GLP1RAs)^8,9^ and TGFβ superfamily inhibitors^10^.

Small molecule inhibitors of DYRK1A as Harmine, INDY, Leucettine, 5-Iodotubericidin (5-IT), GNF and 2-2c, are able to induce human beta cells proliferation *in vitro*, at a rate (or labeling index) of ∼2% as assessed by Ki67, BrdU, or EdU^3-7,11-13^. The addition of a GLP1RA agonist as exendin-4, semaglutide or others, or a TGFβ inhibitor such as LY364947 or GW788388, synergistically increases proliferation to 5-8%^8-11^. Importantly, multiple studies have shown that harmine and other DYRK1A inhibitors, also induce human alpha cells proliferation *in vitro*^3,7,10,14^. Recently, we have shown that human beta cell proliferation could also be induced *in vivo*. Specifically, continuous infusion of harmine and exendin-4 for three months into immunodeficient mice bearing small human islet grafts in the renal capsule, resulted in a 300% (harmine alone) or 700% (harmine + exendin-4) increase in human beta cell mass^15,16^. These remarkable increases in beta cell mass are accompanied by reversal of diabetes, as evident by a return to euglycemia and a marked increase in human insulin secretion^15,16^. Curiously, in contrast to the *in vitro* setting in which the harmine-exendin-4 combination causes an increase in human beta and alpha cell proliferation, proliferation in engrafted human beta cells and alpha cells *in vivo* is modest (∼0.2-0.5%) and can not explain the 300-700% increase in beta cell mass^15,16^. Moreover, despite the dramatic increase in human beta cell mass, there is no accompanying increase in alpha cell mass^15,16^. Collectively, these observations suggest that harmine alone or in combination with exendin-4 may lead to the generation of new beta cells through a mechanism unrelated to replication, such as transdifferentiation from non-beta cell types. Since alpha cells have been repeatedly shown to be activated by harmine and other DYRK1A inhibitors, and along with beta cells, they are the most abundant human islet cell types, and since rodent and human alpha cells are capable of transdifferentiating into beta cells under certain circumstances, we suggest that alpha-to-beta cell transdifferentiation might explain the striking *in vivo* increase in beta cell mass and function occuring in response to harmine^15,16^.

Along these lines, others have previously proposed the existence of endogenous beta cell precursors, progenitors, or stem cells. For example, in the alpha-to-beta transdifferentiation paradigm, Thorel et al. showed that a near complete loss of beta cells in the adult mice leads to conversion of alpha cells to beta cells^17,18^. Importantly, Furuyama et al. have suggested a general plasticity exists among human islet endocrine cells, including alpha cells, which allows them to transdifferentiate from one endocrine cell type to another ^19^. Huising et al. also have identified a subset of urocortin (UCN)-negative, Ins-high, Neurog3-high self-replenishing “virgin” beta cells that appear to be capable of conversion to mature beta cells^20,21^. Kushner et al. have suggested the existence of SOX9^+^/ARX^+^/GCG+/-cells in the human pancreas that might serve as a renewable source of beta cells in people with T1D^22^. Finally, it is also possible that beta cells can arise from other less well characterized islet cell types, including murine acinar cells^23,24^, and PDX1^+^/ALK3^+^/CAII^-^ ductal cells^25,26^ in pancreas samples from humans with T1D and T2D^27^. Collectively, these studies suggest that a variety of normal beta cell precursors may exist in the human pancreas.

Unfortunately, to date, there are no small molecule drugs that induce the generation of *any* human beta cell precursor in numbers that could potentially be clinically relevant. In this study, we suggest that a cluster of “Cycling Alpha Cells”, identified through single cell human islets transcriptomics, may serve as a beta cell regenerative reservoir, and may be readily transdifferentiated into functional human beta cells upon treatment with existing beta cell regenerative drugs of the DYRK1A inhibitor class.

## Methods

### Human islet samples

HIPAA-compliant, de-identified adult human pancreatic islets from four donors (three male and one female) were obtained from Prodo Laboratories (Aliso Viejo, CA), Alberta Diabetes Institute Islet Core (Edmonton, Canada), and Integrated Islet Distribution Program (IIDP). In all cases, informed consent was provided at the institutions where the organs were harvested. The mean age of the donors was 52.75 years (range 47-58 years), and the mean BMI was 30.1kg/m^2^ (range 27.4-32.3kg/m^2^). Purity ranged from 85-90%. Additonal details are provided in Supplemental Table 4.

### Dispersion of human islet cells into single cell suspension and compound treatments

Islets were disassociated with Accutase (Fisher Scientific) as previously described^7-10^. Briefly, islets were centrifuged at 200*g* to obtain pellets of whole islets. Pellets were then washed twice with phosphate-buffer saline (PBS), incubated in Accutase for 15-17 minutes at 37°C, and dispersed into single cell solution by pipette trituration. Islet single cell solution was then pelleted by centrifugation at 700*g* and resuspended in RPMI islet culture medium. Islet single cell solution was then separated into six equal parts each of which consisted of roughly 1000 islet equivalents (IEQs), and was subjected to different compound treatments for 96Hr; DMSO (control), Harmine (10µM), GLP1RA (5nM), LY364947 (3µM), Harmine + GLP1RA (GLP), and Harmine + LY364947 (LY).

### Processing of human islet cells for scRNA-seq

After the completion of 96-hour compound treatment, islets were washed once with PBS and then dispersed with Accutase for 10 minutes at 37°C. Dispersed islets were then collected in RPMI islet culture medium and centrifuged at 700*g* to obtain cell pellets. These pellets were then resuspended in 0.5% bovine serum albumin (BSA) in PBS at 1 x 10^6^ cells/ml. Cells were processed according to Chromium 3’ Gene Expression V3 Kit (10X Genomics) using the manufacturer’s guidelines followed by sequencing on an S1 Novaseq chip (Illumina Inc.) at the Genomics Core of Icahn School of Medicine at Mount Sinai.

### Single cell RNA-seq Analysis, Quality control and preprocessing

FASTQ files were aligned to the GRCh38 human genome reference, filtered, barcoded and UMI counted using Cell Ranger version 7.0.0 (10X Genomics). Empty droplets and substrate ambient RNA remove from count matrices with remove-background function from cellbender (version 0.2.2)^28^. Each dataset was then filtered to retain cells with ≥1000 UMIs, ≥400 expressed genes, and <20% of reads aligned to the mitochondrial genome. UMI counts were then normalized so that each cell had a total of 10,000 UMIs across all genes and log-transformed with a pseudocount of 1 using the “LogNormalize” function in the Seurat package (version 4.0.3, RRID:SCR_016341)^29^. The top 2000 most highly variable genes were identified using the “vst” selection method of “FindVariableFeatures” function and counts were scaled using the “ScaleData” function. Principal component analysis was performed using the top 2000 highly variable features (“RunPCA” function) and the top 30 principal components were used in downstream analysis. Datasets for each donor from DMSO, Harmine, Harmine + GLP and Harmine + LY treatments were integrated by using the “RunHarmony” function in the harmony package (version 0.1.0)^30^ where sample name was used as the group for batch correction. K-Nearest Neighbor graphs were obtained by using the “FindNeighbors” function and the UMAPs were obtained by the “RunUMAP” function. Louvain algorithm was used to cluster cells based on expression similarity. Cell cycle regression of the integrated data is performed by first calculating the cell cycle phases using the CellCycleScoring function of Seurat and then doing the regression by using the G2/M and S phase scores as variables to regress for scaling data prior to harmony integration. Endocrine cells were selected from cell-cycle regressed integrated dataset, and clustering analysis was performed again to generate endocrine-only UMAP. Similarly, cluster 6 (Cycling Alpha concentrated cluster) was isolated from endocrine only integrated data and then clustering analysis was performed again to generate Cycling Alpha UMAP. The resolution was set at 0.4 for the integrated dataset containing all cells and endocrine cells, and at 0.2 for integrated dataset containing only the Cycling Alpha cells for optimal clustering.

### Cell type annotation

Differential markers for each cluster were identified using the Wilcox test (“FindAllMarkers” function) with adjusted p-value < 0.01 and absolute log2 fold change > 0.25, and minimum 10% of cells expressing the gene in both comparison groups using 1000 random cells to represent each cluster. The top up-regulated genes and curated genes from the literature were used to assign cell types to clusters in the all cells integrated dataset, and the expression of marker genes are visualized using DotPlot function. Gene expression signature scores were calculated using AddModuleScore function from Seurat package. To further confirm our cell annotations, first, we performed reference-based mapping using Azimuth R packages (version 0.4.6)^29^ and employed previously annotated human islet cell dataset as reference^31^. Predicted cell annotation was projected on our integrated data UMAP with all cells, and the expression level of annotation markers used by Elgamal et al was visualized by DotPlot^31^. We further performed cell type fractions prediction using Unicell deconvolve (version 0.0.1)^32^. Cell type probability of selected cell types were visualized on UMAP. The initial cell type annotation in the initial integrated dataset was applied to all subsequent datasets and UMAPs.

### Cell abundance analysis

Cell type abundance was visualized using barplot which shows the percentage of each cell type within each treatment condition. To explore changes in abundance of the different cell types between treatment and DMSO samples, MiloR (version 1.2.0)^33^ R package was used to perform differential abundance test in integrated dataset with both endocrine and exocrine cells. Milo graph and neighborhoods were generated using k=20, d=30 and harmony embeddings as dimensionality reduction in integrated dataset of all cells. Cell count differences across samples were normalized, and comparison was carried out using testNhoods function. Neighborhood with cell type fraction <0.7 is considered as ‘mixed’ neighborhood. Results of cell abundance test was visualized using plotDAbeeswarm function in MiloR. After selecting only endocrine cells, percentage of cell type in each treatment was calculated again as number of cells of each cell type in each treatment divided by total number of cells in the treatment. Unstacked barplots were used to visualize the percentage of cell type across treatments, and Dunnett’s test was applied to assess the significance of change in abundance between H, H+GLP and H+LY against DMSO.

### Pseudotime, velocity and PAGA analysis

RNA velocity analysis was performed using the scvelo Python package (version 0.2.5) on integrated endocrine cells data^34^. The unspliced and spliced count matrices were generated from Cell Ranger output using the run10x function from velocyto package (version 0.17.17)^35^. For all-sample-integrated endocrine cell dataset, Seurat object was converted into anndata (version 0.8.0)^36^ and merged with the unspliced and spliced count matrices using scanpy (version 1.9.3)^37^. Moments for velocity estimation were calculated using 30 principal components and 30 neighbors. RNA velocity was estimated using stochastic model with default parameters using scvelo. Cell type labels from the Harmony integrated endocrine cell dataset were projected to the all-sample-integrated dataset, and velocity streams were visualized. To explore the connectivity between cell clusters, partition-based graph abstraction (PAGA) was calculated based on velocity results using the PAGA wrapped in scvelo^38^. pyGPCAA wrapped in CellRank was utilized to find initial and final state clusters in integrated all cell dataset, using the abovementioned velocity information (version 1.5.1)^39^. To infer progression of cells through geodesic distance along the graph, diffusion pseudo-time was calculated using the dpt tool from scanpy, with a cell from the initial cluster selected by CellRank as the root cell.

### Gene Regulatory Network Analysis

To infer gene regulatory networks in each cluster, pySCENIC (v0.12.1, RRID: SCR_017247) was applied to the integrated endocrine cells dataset^40^. The list of transcription factors for hg38 human genome was downloaded from https://resources.aertslab.org/cistarget/tf_lists/ and was used for querying transcription factors of the pySCENIC analysis. Motif annotation and the cisTarget datasets were obtained from https://resources.aertslab.org/cistarget/motif2tf/ and https://resources.aertslab.org/cistarget/databases/homo_sapiens/hg38/refseq_r80/mc_v10_clust/gene_based/ respectively. Briefly, co-expression of genes and transcription factors was first inferred from gene expression matrix using grn function and the co-expressed genes and transcription factors together define gene regulons. The regulons were further refined by finding only cis-regulatory target of each transcription factor using ctx function for motif discovery. Finally, the cellular enrichment of each refined regulon was calculated using aucell function from pySCENIC. In each annotated endocrine cell type cluster, regulon specificity score based on the Jensen-Shannon divergence was calculated to evaluate the activity of each regulon. Regulon specificity score was visualized using clustermap function from seaborn (v 0.12.1, RRID: SCR_018132)^41^.

### Code Availability

All analysis is performed by publicly available packages and no custom method is implemented. All the packages used have been cited throughout the manuscript.

### Data Availability

All transcripomic data are available in dbGaP under accession # TBD, and via reasonable request from the corresponding author. All supplemental tables can be accessed via the following link: https://www.dropbox.com/scl/fo/3i21g61kd3s6qvmkza4nw/h?rlkey=8ly7yucvzryrje98dq8be2rdp&dl=0

## Results

### Single Cell transcriptomic analysis of human islets

Human islets from four otherwise healty adult islet donors were treated with 10μM harmine alone, 10μM harmine together with 5nM of the GLP1R agonist exendin-4, or 10μM harmine together with 3μM of the TGFb inhibitor, LY364947 (LY) for 96 hours and subjected to single cell transcripomics (**Suppl. Fig. 1A**). We identified 21 unique cell type clusters and confirmed the prescence of all previously identified cell types in human islets^31,42^ (**Fig. 1A**; **Suppl. Fig. 1B**). All annotated cell types were present in all four islet donors, minimizing residual batch effects driven by individual donors as data was integrated and batch-corrected using the ‘harmony’ tool (**Suppl. Fig. 1C**). We also defined the cell cycle stage for each cell type (**Suppl. Fig. 1D**). Finally, we also demonstrated the presence of hormone-producing cells by generating feature plots showing the expression level of islet cell type hormones, insulin (*INS)*, glucagon (*GCG)*, somatostatin (*SST*) and pancreatic polypeptide (*PPY*) (**Suppl. Fig.1E**). Collectively, these findings indicate the islet cells and subtypes identified here correspond closely to those identified in prior human islet scRNAseq and snRNAseq datasets^31,42^.

**Figure 1.**
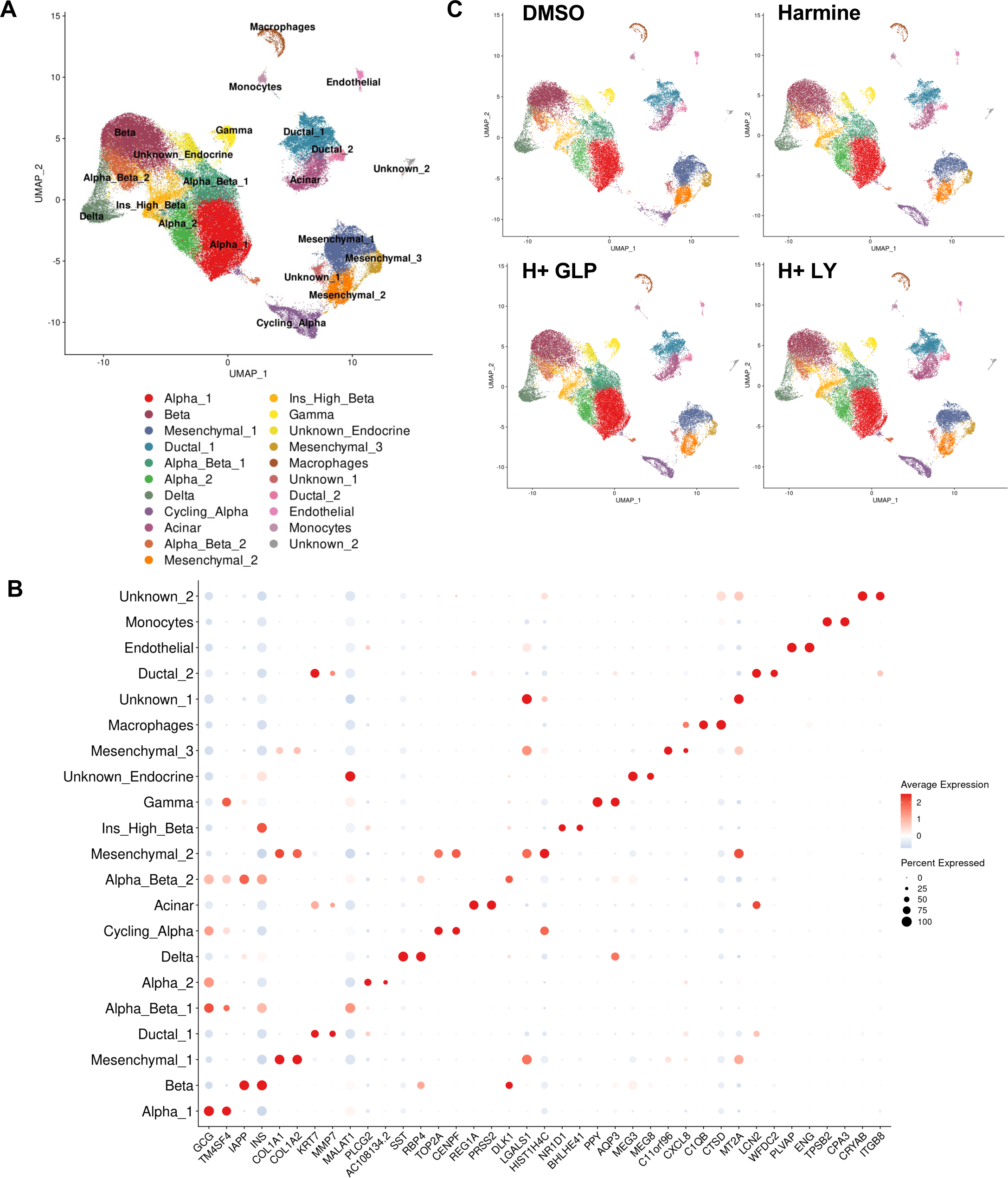
Regenerative Drug Treatment on Human Islets Increases the Abundance of Cycling Alpha Cell Population. **A.** UMAP plot showing clustering of single cell RNA-seq profiles from 109,881 cells from the integrated dataset. Clusters are labeled based on cell type specific marker gene expression. **B.** Dot plot showing average expression level and percent expressing cells of the top two upregulated genes in each cluster compared to all other clusters. **C.** Split UMAP plots showing single cell RNA-seq profiles from each treatment group.

### Dataset annotations are congruent with the HPAP dataset and the deep learning model, Unicell

To further benchmark our annotations with previously published datasets, we first projected our integrated dataset onto the HPAP human pancreas reference dataset from the PANC-DB website using the publicly available Azimuth tool^29,31,43^. This provided high confidence alignment of multiple cell populations (**Suppl. Fig. 2A,B**) with a high mapping and prediction scores (**Suppl. Fig. 2C,D**). Furthermore, we also predicted cell types in our integrated dataset using unsupervised deep learning method called UniCell^32^. UniCell analysis confirmed our previous cell type annotations for endocrine, beta, alpha, ductal, acinar and connective tissue cells (**Suppl. Fig. 3A-F**). Intriguingly, a small fraction of the Cycling Cells was annotated as a connective tissue cells (**Suppl. Fig. 3F**).

### Cycling Alpha Cells uniquely fulfill characteristics for the regenerative drug-responsive target cells

Among endocrine cell clusters, two Beta (labelled as “Beta” and “Ins High Beta”), two Alpha-Beta and two Alpha cell clusters were identified (**Fig. 1A**). Between the two alpha cell clusters, Alpha1 appears to represent more mature or differentiated alpha cells, showing higher experssion of *GCG*, *TM4SF4*, *CRYBA2* and *CHGB* genes (**Supp.** Fig.1B). Among the two Alpha-Beta clusters, Alpha-Beta1 selectively expressed alpha cell identity genes as well as *INS* and *PCSK2*. In contrast Alpha-Beta2 cluster expressed a broader group of alpha and beta cell identity genes (**Suppl. Fig. 1B**). With respect to cell cycle activity, two distinct cell types showed high levels of proliferating cell markers such as *MKI67*, *TOP2A*, *CENPF* and *NUSAP1* expression (**Suppl. Fig. 1B**). Of these two cell types, ‘Cycling Alpha’ cells expressed mainly alpha cell identity genes (*GCG*, *CHGB* and *CRYBA2*) as well as the insulin granule-associated gene, *SCG2* (**Suppl. Fig. 1B**) suggesting the presence of a proliferating endocrine cell cluster present in all four donors.

### Regenerative drug treatment leads to expansion of Cycling Alpha Cells

We next compared the proportions of each cell type across samples (**Fig. 2A**). To correct for the cell number variability in different donors, we assessed the proportion of each given cell type by normalizing it to the total cell number. As previously reported, alpha cells represent the largest proportion of islet cells, followed by beta cells. Cycling Alpha Cells were the eighth largest population. Notably, treatment with human regenerative drugs consistently increased the percentage of Cycling Alpha Cells as compared to control treatment (**Fig. 2A**).

**Figure 2.**
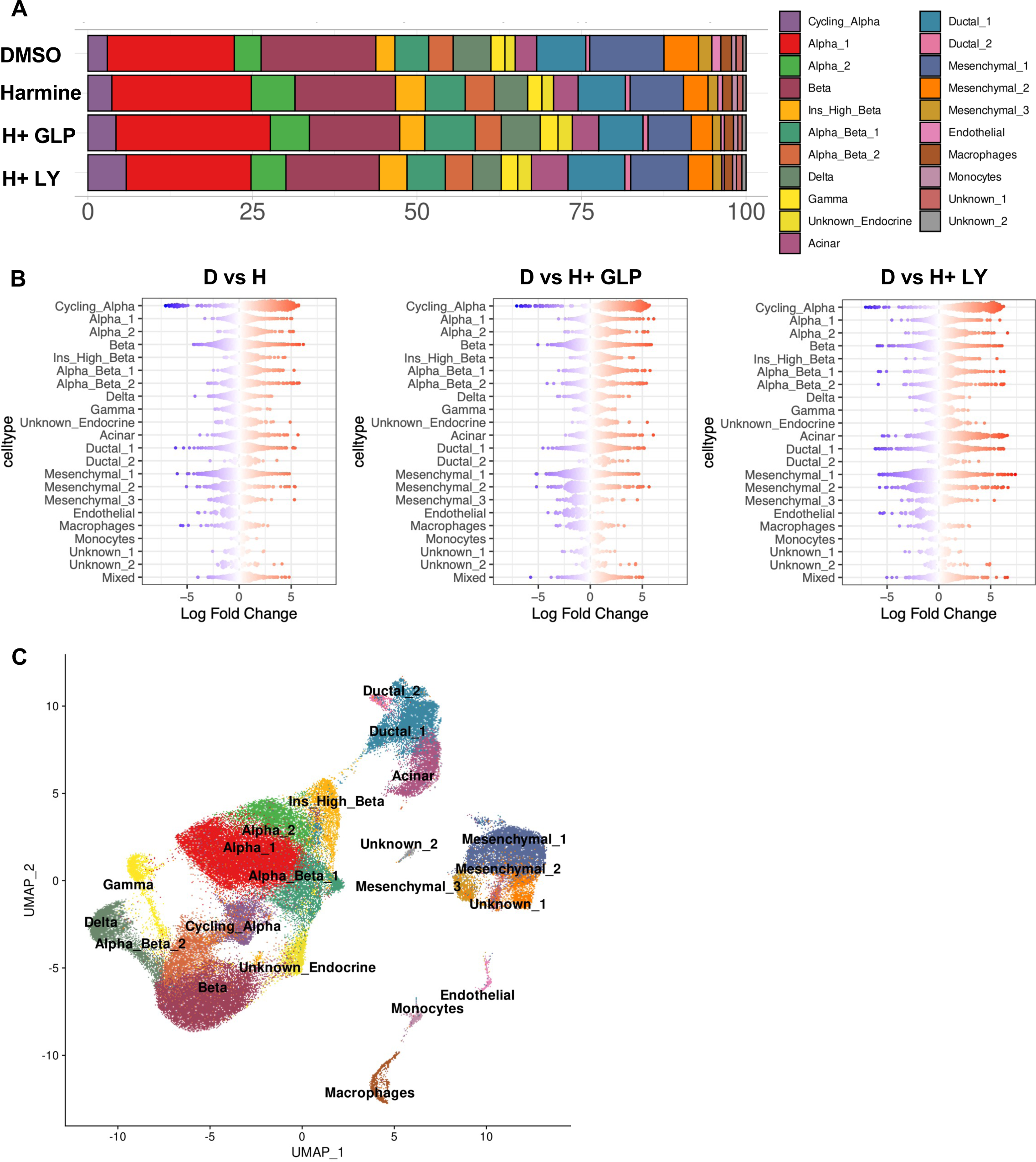
Cell Cycle Regression Places Cycling Alpha Cells in the Endocrine Subset. Cycling Alpha Cell Cluster Is the Only Cluster that Becomes More Abundant with Regenerative Drug Treatment. **A.** Proportion of cells from each cell type, grouped by treatment. **B**. Differential abundance of islet cell types in different contrast groups, assessed by Milo statistical framework. **C.** UMAP plot depicting unsupervised clustering of single cell RNA-seq profiles after cell cycle regression. Cell are colored by initial annotation.

We also noted a possible increase in the percentage of other cell types, exemplified by acinar cells, in **Fig. 2A**. To statistically quantify the changes in cell abundances, we used MiloR^33^ to perform differential abundance testing by assigning cells to partially overlapping neighborhoods (**Fig. 2B**). Among all cell types, Cycling Alpha cluster demonstrated the strongest increase in abundance after drug treatment, underlining the impact of regenerative drug treatment on Cycling Alpha Cells (**Fig. 2B**). Taken together, these results indicate that a four-day treatment of human islets with beta cell regenerative drugs substantially increased the abundance of this previously understudied Cycling Alpha Cell cluster.

As seen in **Figs. 1A** and **C**, Cycling Alpha Cells, which express high levels of alpha cell markers, cluster more closely to mesenchymal cells than to endocrine cells. This could potentially be explained by the effect of the cell cycle phase rather than the cell type. To explore this possibility, we repeated the identical unsupervised clustering, but we first regressed the cell cycle genes to eliminate their effect on clustering. This approach clearly showed that after cell cycle regression, Cycling Alpha Cells now clustered among the other endocrine cell populations, and connected to alpha, beta and alpha-beta cell clusters (**Fig. 2C**).

### Cycling Alpha Cells expand and increase beta cell identity marker expression following regenerative drug treatment

To better delineate the sub-populations and associated molecular signatures in the Cycling Alpha cluster, we removed all cell types except those annotated as endocrine cells from the cell-cycle regressed integrated dataset, and performed dimensional reduction and unsupervised clustering again to focus our analysis on endocrine compartment (**Suppl. Fig. 4A**). This yielded twelve separate cell clusters at resolution 0.4, ten of which perfectly matched our previous cell identity annotations. The two new clusters reflected: the division of Alpha1 and Alpha-Beta1 into three clusters (clusters 0, 2 and 10); and, the sub-division of the Cycling Alpha cluster into two sub-clusters (clusters 6 and 11) with distinct molecular signatures. Remarkably, cluster 11 was separated from all endocrine clusters despite being clustered together with other Cycling Alpha Cells before cell-cycle regression and showing high expression of proliferation markers. Instead, cluster 11 was enriched for high levels of mesenchymal marker genes, such as COL1A1, IGFBP5 among others (**Suppl. Fig. 4B, C**). Moreover, regenerative drug treatment led to attenuation or disappearance of this cluster in three of four islet donors when compared to DMSO treatment (**Suppl. Fig. 5**). Accordingly, due to its clear mesenchymal character, we excluded cluster 11 from the endocrine cell populations, and repeated previous unsupervised clustering and analyses.

This new analysis yielded eleven separate clusters at resolution 0.4, all of which, as expected, were endocrine cells (**Fig. 3A, B**). Projection of cell type annotations on these endocrine cells indicated a clear overlap between our previous annotations and the new clustering (**Fig. 3B**) with the exception, as expected, the absence of a mesenchymal cell cluster in this dataset (**Fig. 3C, Suppl. Fig. 6A, B**). We also verified that the abundance of Cycling Alpha Cells was significantly higher with regenerative drug treatment vs. control treatment (**Fig. 3D**). We next investigated differentially expressed cluster-specific genes, and found that 878 genes were upregulated in the Cycling Alpha cluster (cluster 6) vs. the rest of the clusters (**Suppl. Table 1**). Hallmark Gene Set enrichment analysis on these genes demonstrated that cell cycle-related pathways, exemplified by ‘E2F Targets’ and ‘G2M Checkpoint’ were highly upregulated in Cycling Alpha Cells (**Fig. 3E**). In addition, we found that DREAM complex transcription factors are highly activated in Cycling Alpha Cells, based on the gene regulatory network analysis performed by pySCENIC and on the enrichment of DREAM complex members gene expression in Cycling Alpha Cells (**Suppl. Fig. 7**). Overall, in addition to the increase in cell abundance (**Fig. 3F**), the most important observation was that while cells present in the Cycling Alpha cluster under basal conditions had a predominant alpha cell phenotype, Cycling Alpha Cells appeared to acquire a beta-like phenotype in response to regenerative drug treatment (**Fig. 3G**).

**Figure 3.**
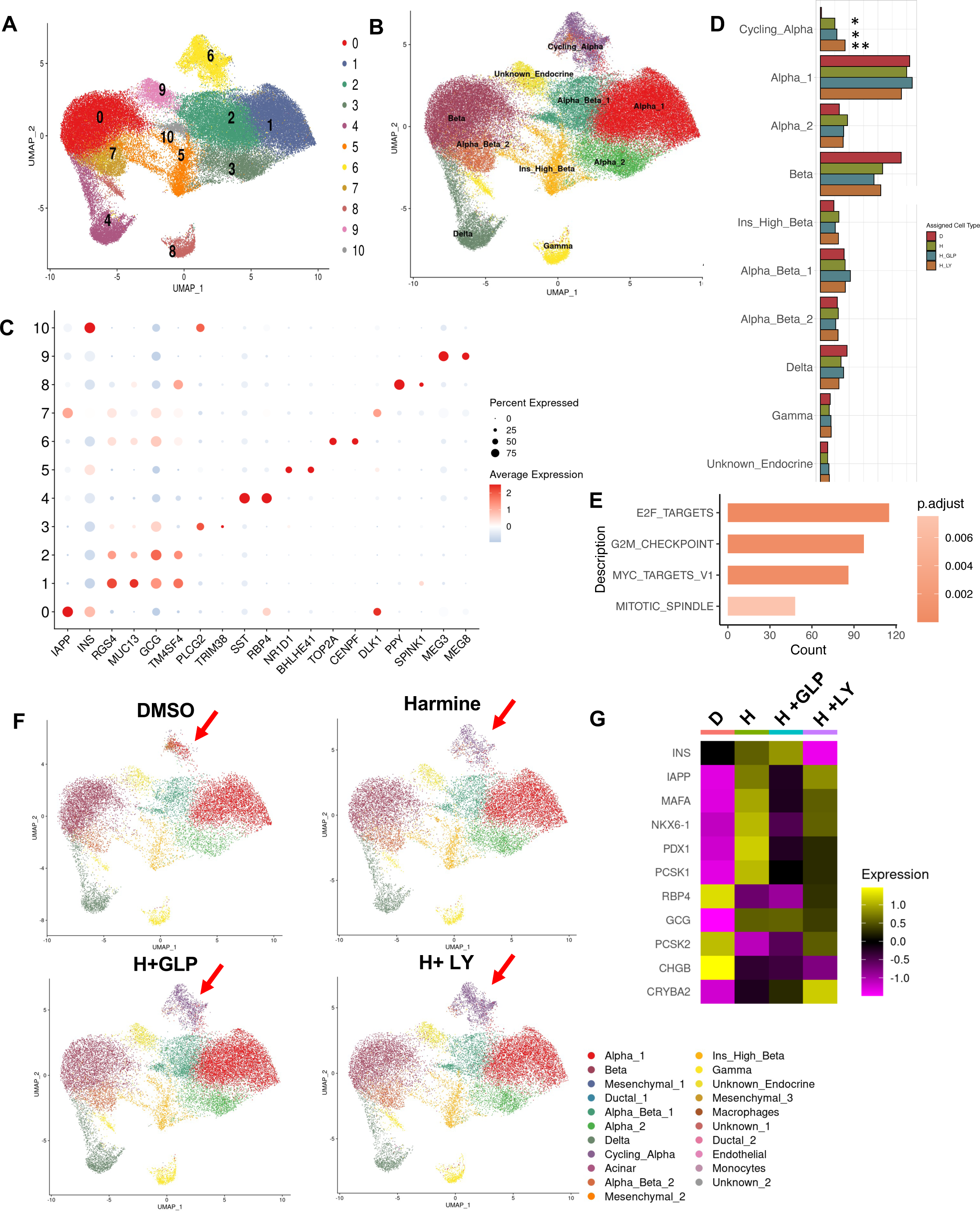
Apparent Conversion of Cycling Alpha Cells into Beta Cells. **A.** UMAP plot showing unsupervised clustering of single cell RNA-seq profiles of only endocrine cells after the removal of subcluster 11. **B.** Same UMAP plot as in A, depicting cell annotation projections. **C.** Dot plot showing average expression level and percent expressing cells of the top two upregulated genes in each cluster. **D.** Proportion of cells from each cell type, grouped by treatment. Note that Cycling Alpha cells are the only cell type with a significant increase after regenerative drug treatment. **E.** Hallmark Gene Set Enrichment analysis on the upregulated genes in Cycling Alpha Cells (cluster 6) vs rest of the endocrine cell types. **F.** Split UMAP plots showing single cell RNA-seq profiles from each treatment group. Arrows depict the Cycling Alpha Cluster. **G.** A heatmap showing the expression level of selected alpha- and beta-cell marker genes under drug treatment in Cycling Alpha Cells. (Significance was calculated by Dunnett’s test. * means p<0.005, ** means p<0.0001)

Intrigued by this observation, we further sub-clustered the Cycling Alpha Cells (cluster 6) from the integrated dataset shown in **Fig. 3A**, and demonstrated that it yielded a total of three subclusters at resolution 0.2 (**Fig. 4A**). Exploring the cell cycle phase of the cells in Cycling Alpha cluster, we found that cells pertaining to all three cell cycle phases were present (**Fig. 4B**). Moreover, we assessed the cell identity annotations by assigning alpha cell, beta cell and cycling module scores. Our results demonstrated that while alpha cell module score was high across the board, cells with a higher alpha cell score were found in cluster 0, and cells with a high beta cell module score was restricted to cluster 1. In addition, cycling module score was high in cluster 2 and partly in cluster 0, but not in cluster 1 (**Fig. 4C**). We found that cluster 2, as well as part of cluster 0 with a high cycling module score, contained the fewest cells in the DMSO condition, and these two clusters expanded with each drug treatment, indicating that regenerative drug treatment induces an important expansion among Cycling Alpha Cells (**Suppl. Fig. 8**). We further confirmed cell identities by assaying differentially regulated genes in each cluster (**Fig. 4D, Suppl. Table 2**). Hallmark Gene Set enrichment analysis on the differentially upregulated cluster genes also revealed that cluster 2 had high enrichment for cell cycle related terms while beta cell related terms were highly enriched in cluster 1 (**Fig. 4E**). Since cluster 1 had the highest beta cell module score and did not display proliferative capacity based on the differentially expressed genes as well as the cycling module score, we hypothesize that the cells present in other clusters may be capable of trans-differentiating into beta cells. Collectively, this analysis suggests that Cycling Alpha Cells can be encouraged to acquire a beta cell phenotype with regenerative drug treatment.

**Figure 4.**
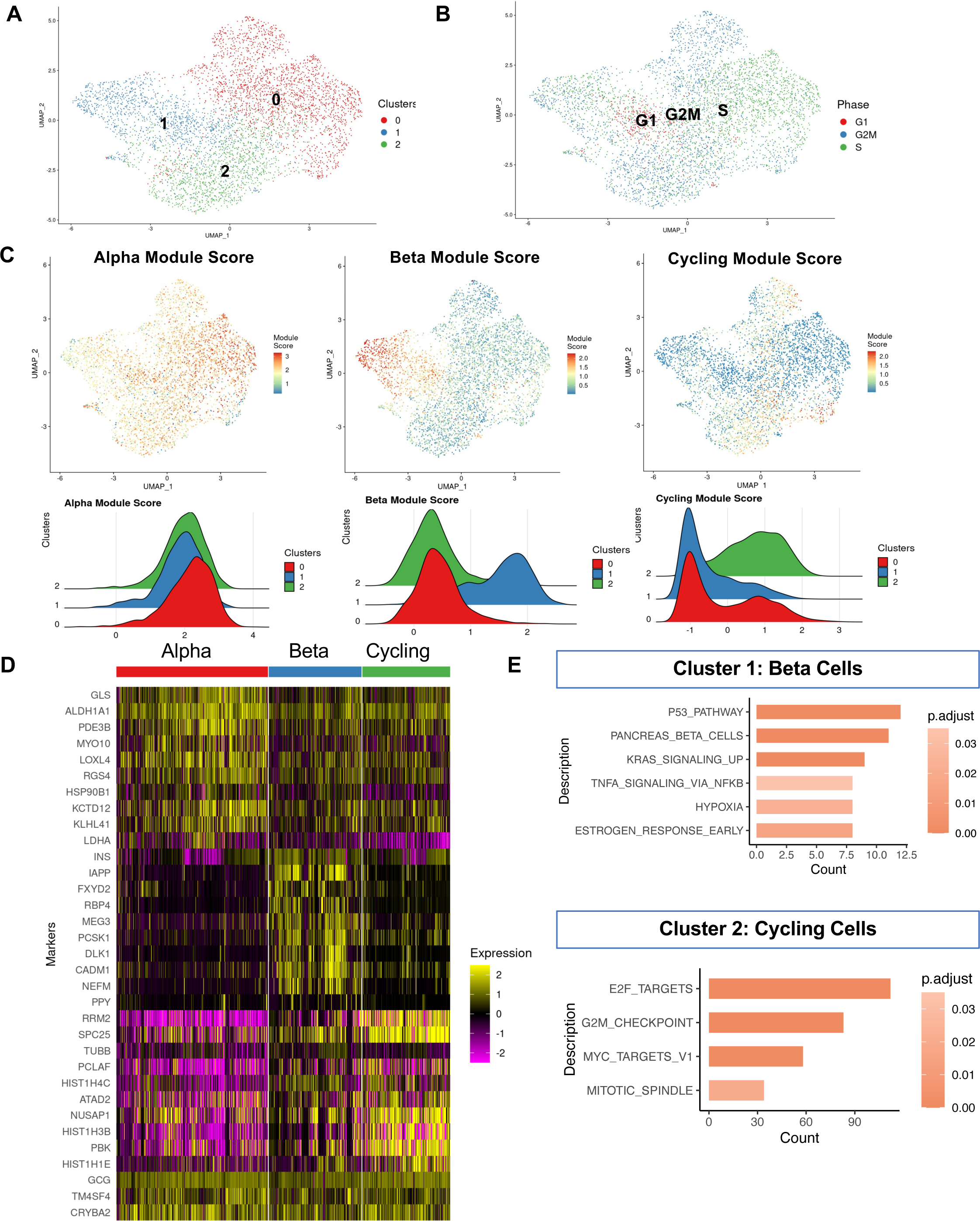
Cycling Alpha Cell Cluster in the Integrated Dataset Is Composed of Alpha, Beta and Cycling Cells. **A.** UMAP plot showing unsupervised clustering of single cell RNA-seq profiles of only cluster 6 (cycling cells and endocrine cells clustered closely together with cycling cells) present in the endocrine cell population. **B.** UMAP plot showing same cells as in A, labeled based on cell cycle phase. **C.** UMAP plots showing cells colored based on Alpha, Beta and Cycling Module Scores. **D.** Heatmap depicting top ten upregulated genes in each of the sub-clusters in the cycling endocrine cells. Last three genes (GCG, TM4SF4 and CRYBA2) are manually included as alpha cell markers. **E.** Hallmark Gene Set Enrichment analysis on subcluster1 and subcluster2 in the cycling endocrine cells.

### Trajectory inference suggests that Cycling Alpha Cells can transdifferentiate into beta cells in response to regenerative drug treatment

To test the hypothesis that a subpopulation of Cycling Alpha Cells may transdifferentiate into beta cells or other endocrine cells, we performed RNA velocity analysis to quantify cellular transitions using the integrated endocrine cell dataset from all four donors. The results suggest that Cycling Alpha Cells are indeed capable of differentiating into Alpha-Beta2-type cells, which in turn can give rise to beta cells (**Fig. 5A**). Importantly, through coarse-grains cell-cell transition matrix onto the macro-state level, CellRank identified the Cycling Alpha Cluster as the initial state (**Fig. 5B**). In line with the velocity results, diffusion pseudotime analysis also predicted Cycling Alpha Cells can transit into Alpha-Beta cells and beta cells when Cycling Alpha Cell were selected as initial cluster (**Fig. 5B**). These findings collectively suggest that a Cycling Alpha to Alpha-Beta2 to Beta cell transdifferentiation axis is present in human islets subjected to regenerative drug therapy. To gain a deeper understanding of the molecular pathways involved in these presumed trans-differentiation events, we also identified the genes that are expressed along the pseudotime axis (**Fig. 5C, Suppl. Table 3**). Taken together, trajectory inference by RNA velocity and pseudo-temporal ordering, strongly suggest that Cycling Alpha Cells have the potential to transdifferentiate into beta cells in response to regenerative treatment.

**Figure 5.**
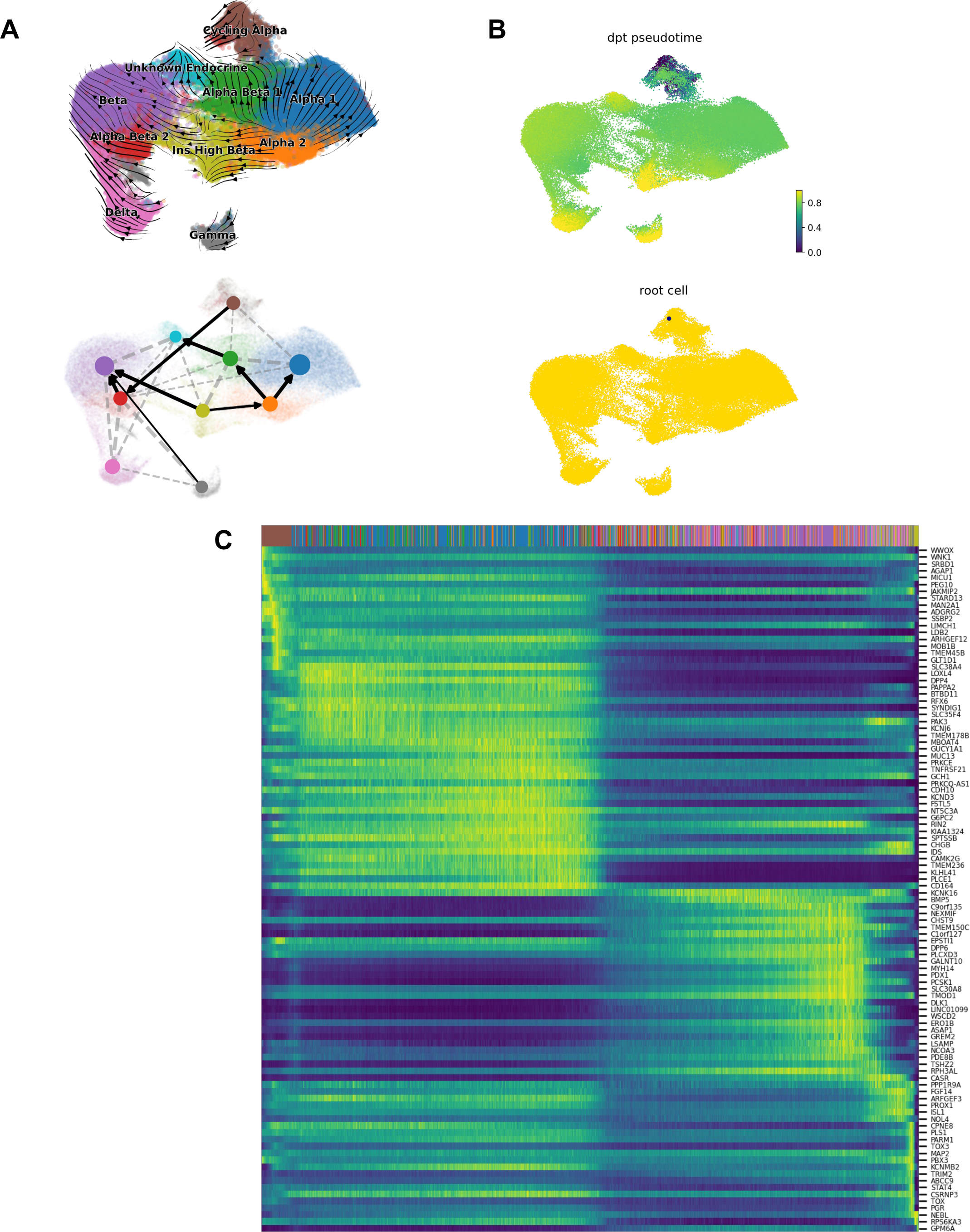
Velocity and Pseudo-Time Analyses Reveal Cell Fate Determination after Regenerative Drug Treatment on Human Islets. **A.** Velocity analysis and Partition-based Graph Abstraction (PAGA) on endocrine cells. **B.** Pseudotime trajectory analysis on endocrine cells. Note that Cycling Alpha Cells are picked as root cells by the GPCAA algorithm. **C.** Heatmap depicting genes expressed along the pseudo-time axis.

## Discussion

DYRK1A inhibitors are able to drive human beta cell proliferation *in vitro and in vivo*, an effect that can be further augmented by combined treatment with number of GLP1 receptor agonists, or TGFβ superfamily antagonists, as previously shown^3-10,44,45^. For example, the DYRK1A inhibitor, harmine, increases beta cell mass in human islets transplanted into immunodeficient mice by 300% over three months, and the beta cell mass is further increased to 700% by the addition of the GLP1RA, exendin-4 or exenatide^15,16^. Harmine treatment also leads to enhanced beta cell function *in vitro* and *in vivo*, evidenced by enhanced glucose-stimulated insulin secretion, enhanced expression of key beta cell transcription factors and functional markers (e.g., PDX1, MAFA, NKX6.1, SLC2A2, and INS itself), and a rapid (days) return to euglycemia in diabetes models^3,7-10,15,16^. Yet, one conundrum remains unexplained: in contrast to the remarkable 3-8% increases in beta cell proliferation observed in cultured beta cells *in vitro*, as assessed by Ki67 or BrdU labeling, beta cell proliferation assessed using Ki67 labeling in transplanted human islets *in vivo* is modest (0.3-0.6%), and cannot explain the 300-700% increases in human beta cell mass that occur over three months *in vivo*^15,16^. Here, we report that beta cell regenerative drug treatment of human islets leads to a substantial increase in the abundance of the islet cell subtype called “Cycling Alpha Cells”, suggesting that transdifferentiation of these Cycling Alpha Cells into beta cells is an important contributor to the dramatic increase in new beta cell mass, appearing in human islets in response to treatment with DYRK1A inhibitors.

This study is the first to report single cell transcriptomics data for regenerative drug-treated human islets. From a data quality and analysis standpoint, the datasets in this report, the islet cell subtypes, clusters and annotations correlate closely with datasets from multiple prior studies^31,42,46-53^, suggesting that cell-type identification is accurate and in line with prior studies. In particular, the “Cycling Alpha Cell” cluster has been reported in multiple prior studies^31,46,54-59^. However, the representation of this cluster among other endocrine cells is small unless treated with regenerative drugs. Critically, deeper analysis of this cluster in human islets treated with vehicle, harmine alone or with GLP1 or the TGFβ inhibitor LY364947 (**Fig. 4A-C**), supports the concept that subsets of these Cycling Alpha Cells acquire beta cell characteristics. In addition, RNA velocity and pseudotime analysis of the Cycling Alpha Cells points to a transdifferentiation event, converting Cycling Alpha to Alpha-Beta cells, which then convert into Beta cells (**Fig. 5A**). Alpha-to-beta transdifferentiation is further corroborated by the significant changes observed in the transcriptomic profile of Cycling Alpha Cells upon regenerative drug treatment (**Fig. 3G**, **Suppl. Table 1**). Furthermore, the most significantly upregulated genes in the Cycling Alpha Cells were related to ‘E2F targets’, ‘Myc targets’, ‘G2M checkpoint’ and ‘Mitotic Spindle’ pathways, all of which are consistent with cell cycle initiation and harmine-mediated inhibition of the DREAM complex^60^ (**Fig. 3E**). Taken together, we propose that a novel mechanism for the large and remarkable increase in human beta cell mass observed in response to these drugs *in vivo* is likely a conversion or transdifferentiation of Cycling Alpha Cell precursors into beta cells.

Spontaneous alpha-to-beta cell transdifferentiation has not previously been reported to occur in response to any type of beta cell regenerative drug therapy. However, previous studies reported alpha-to-beta cell transdifferentiation, as described above in the Introduction. As additional examples, Collombat et al. showed that overexpressing *Pax4* in mouse alpha cells leads to beta cell conversion and reverses hyperglycemia in juvenile streptozotocin (STZ)-treated mice^61^. In another study, Zhang et al. demonstrated that adenoviral delivery of *Pax4* in an alpha cell line suppressed glucagon and induced insulin synthesis^62^. In a separate study, inactivation of *Arx* and *Dnmt1* in adult mouse pancreatic alpha cells led to conversion of alpha cells to beta cells with the ability to secrete insulin upon glucose stimulation, although the insulin secretory capacity of these cells was less robust than from mature beta cells^63^. Pharmacological attempts at alpha-to-beta cell conversion include artemisinins that stimulate GABA signaling, translocating ARX to the cytoplasm and promoting its degradation^64,65^, although these findings are controversial^66,67^. Xiao et al. demonstrated that adeno-associated virus (AAV)-delivery of *Pdx1* and *Mafa* into the murine pancreatic duct or to human pseudoislets leads to the conversion of alpha to beta cells^68^. Importantly, Thorel et al. showed that near complete loss of beta cells leads to alpha-to-beta cell conversion in diabetic mice, confirmed using lineage tracing^17,19^. It has also been suggested to occur in FACS-sorted human alpha cells transplanted into mouse models^19^.

While the possibility of human alpha-to-beta cell transdifferentiation has appeal from the current report as well as from prior mouse studies, its occurrence is impossible to unequivocally document at present, because appropriate lineage tracing technologies do not yet exist for human alpha cells. More specifically, although in theory alpha cell lineage tracing tools used in mice could be transferred to humans, there is currently no human alpha cell-specific promoter, an essential tool for such studies. Indeed, multiple human alpha cell promoters (e.g., glucagon, ARX, TM4SF4, GC) that are effective in mice, fail to mark human alpha cells with efficiency or specificity. Similarly, FACS or magnetic bead sorting of human alpha cells can be performed, but yields of alpha cells are poor, and impure, contaminated by other cell types. Thus, drug-induced human alpha-to-beta cell transdifferentiation remains to be proven in unequivocal experimental terms.

In addition to alpha-to-beta conversion, additional cell type conversion scenarios may occur with regenerative drug treatment of islets. For example, conversion of the Alpha2 cluster to other alpha and beta cell types, or Insulin-High beta cells to beta cells, or Unknown Endocrine cells or Alpha-Beta and even Delta cells to beta cells all appear to occur, but as the cell abundances of these clusters didn’t show clear difference between drug treatment and DMSO, we focused on Cycling Alpha Cells (**Fig. 5A**). Thus, future studies will undoubtedly shed light on the lineage dynamics of the endocrine cell types in the presence of regenerative drugs. Yet, among these, Cycling Alpha Cells was predicted to be the ‘root’ cells from the transition model constructed by Generalized Perron Cluster Cluster Analysis (GPCAA) using RNA expression and velocity data, and a Cycling Alpha to Alpha-Beta2 to Beta cell trans-differentiation axis is readily detectable (**Fig. 5B**). Along the pseudotime trajectory, Delta, Gamma and Insulin-High beta cells appear to be in the very end, which may suggest that Cycling Alpha Cells also have the potential to convert into these cells. Further studies are needed to confirm these findings in human islets.

Several prior authors, including ourselves, have suggested that DYRK1A inhibitors induce human alpha cells to proliferate in human islets *in vitro*^3-7,45^. In retrospect, taken together with our current findings, we suggest that DYRK1A inhibitors induce proliferation in the alpha cell precursors of eventual beta cells. This scenario would explain Ki67 or EdU labeling of alpha and beta cell populations that we and others have reported^7-9,15,59,60^, as well as their conversion to beta cells with an ultimate expansion of beta cell, but not alpha cell numbers^15^. Importantly, if this “alpha cell progenitor scenario” is correct, attempts to develop diabetes regenerative drug-targeting approaches directing drugs exclusively to mature beta cells may be mis-directed.

Others have previously suggested the existence of additional putative human beta cell precursor cell types, as described in the Introduction^20-27^. How Cycling Alpha Cells might compare in retrospect to Virgin Beta Cells, SMAD7^+^, BMP7^+^, SOX9^+^, NGN3^+^, MAFA^+^, PDX1^+^, CK19^+^ or other putative beta cell progenitors described by others remains uncertain. None of the markers listed above is differentially expressed in the Cycling Alpha Cell populations upon regenerative drug treatment (data not shown).

We were surprised not to observe a population of proliferating beta cells following DYRK1A inhibitor treatment. It is possible that Ki67^+^ beta cells observed in islets by ourselves and others were bihormonal alpha-beta cells. It is also possible that the beta cells in the four human islet preparations used were less responsive than the average human islet donors, although the donor islets appeared to be of high quality. In future studies, it will be important to assess Ki67 immunolabeling and assessment for bihormonal alpha-beta cells.

This study has important limitations, most of which reflect technical limitations in human islet research. First, we included only four human islet donors in this study, yet human islet preparations are highly variable in composition and quality^69,70^. On the other hand, we paid great attention to the selection of high-quality islets and included islets from donors similar in age, sex and BMI (**Suppl. Fig 1A, Suppl. Table 4**). Reassuringly, our cell type composition closely matched much larger donor sets from human islet scRNAseq and snRNAseq^31,46^. More importantly, the results were highly reproducible, with the key findings being observed in islets from each of the four individual donors (**Suppl. Fig. 1C and 5**).

A second limitation is that these studies were performed on human islets *in vitro.* Alpha and beta cell proliferation rates are substantially higher (∼2-8% Ki67 or BrdU labeling) *in vitro* than they are *in vivo* (0.2-0.6% Ki67 labeling), yet *in vivo* is where a 300-700% increase in beta cell mass is observed^15,16^. Thus, we believe that alpha cell transdifferentiation is even more important in regenerative drug-treated human islet grafts *in vivo*. Accordingly, performance of parallel studies in human islet grafts *in vivo* will be critical in the future.

A third limitation is that the study necessarily lacks human alpha cell lineage tracing confirmation as noted above. This must be performed if and when such technologies become available. Along these same lines, in our experimental design we selected a single time point (96 hours) for regenerative drug treatment. This makes it hard to predict the precise lineage trajectory of proliferating Cycling Alpha Cells. Clearly, drug treatment studies at multiple time points (e.g., 3, 6, 9, 12 days) may be of interest, but human islets survive and retain phenotype poorly over more than a few days. In this regard, Title et al reported that reaggregated pseudoislets can be cultured and subjected to DYRK1A inhibitor treatment for up to 15 days^71^. These reaggregated pseudoislet models hold promise for longer term human islet studies. Nonetheless, with the current absence of reliable alpha cell lineage tracing tools, such longer term studies still cannot provide unequivocal proof of lineage of origin.

A fourth limitation is that while single cell RNAseq studies evidently provide a clear insight into the transcriptomic profiles, a better understanding of transcriptional regulation and developmental lineage likely can be achieved through study of the chromatin dynamics of the same cell types and sub-types identified in a given dataset. Although our results suggest that transcriptional regulators are important for the proliferative effects of DYRK1A inhibitors (**Suppl. Fig. 7**), these findings are computational, and require experimental validation. A more complete and comprehensive understanding of the mechanisms that control transcriptional regulation will require multi-ome studies that combine scRNA-Seq and scATAC-Seq methodologies.

Finally, how fast are these “Cycling Alpha Cells” cycling? Or are they cycling at all? They are annotated as “cycling” because they express certain cell cycle genes and are annotated as being in G1, S or G2M cell cycle phase: it is telling that they exist in presumably quiescent, vehicle-treated islets as well (**Fig. 3F**). If they are really cycling, how “fast” they might be cycling, and how that might compare to prior reports using *in vitro* and *in vivo* Ki67 or BrdU or EdU measurements remains unknown.

Taken together, these studies suggest a novel mechanism of action through which DYRK1A inhibitors are able to expand human beta cell numbers. We suggest that they act via a combination of alpha cell precursor replication followed by alpha-to-beta cell transdifferentiation. If and when critical alpha cell lineage tracing tools become available, future studies will be required to confirm or exclude DYRK1A inhibitor-induced alpha-to-beta transdifferentiation. Finally, since this alpha-to-beta mechanism eliminates, in theory, the need for residual beta cells in order for the beta cell regenerative drugs to reverse diabetes, beta cell regenerative DYRK1A inhibitor drugs may be feasible even in people with T1D and T2D with few or no residual beta cells. Finally, if alpha cells are the principal target for “beta cell regenerative drug therapy”, it raises the question as to whether beta cell drug targeting strategies are necessary for beta cell regeneration, or instead are mis-directed. These issues and limitations should be the focus of future studies.

## Acknowledgements

The authors wish to thank Bonnie and Joel Bergstein, Lonnie and Thomas Schwartz and Martha and Fred Farkouh families for their constant support of this research. We also thank the NIDDK-supported Human Islet and Adenovirus Core (HIAC) of the Einstein-Sinai Diabetes Research Center (ES-DRC), the NIDDK Integrated Islet Distribution Program (IIDP) and Prodo Laboratories for supplying human organ donor islets. We also thank the Genomics Core at the Icahn School of Medicine. This work was supported NIH Grants K-01 DK128378, P-30 DK 020541, R-01 DK130300, R-01 DK126450, R-01 DK125285, R01 DK105015, R-01 DK129196, by the Bioinformatics for Next Generation Sequencing (BiNGS) shared resource facility within the Tisch Cancer Institute at the Icahn School of Medicine at Mount Sinai, which is partially supported by NIH grant P30CA196521, by the computational resources and staff expertise provided by Scientific Computing at the Icahn School of Medicine at Mount Sinai, by the Clinical and Translational Science Awards (CTSA) grant UL1TR004419 from the National Center for Advancing Translational Sciences, and by the NIH Office of Research Infrastructure grant S10 OD 026880.

## Author Contributions

E.K., P.W., and O.W. performed experiments. E.K., X.W., S.C., D.H., L.L., R.K., G.L. and C.A. analyzed data. E.K., C.A., A.G.O., D.K.S., and A.F.S. conceived of the studies. E.K. and A.F.S wrote the manuscript with comments from X.W., S.C., and D.H.

---

**Supplementary Figure 1.**
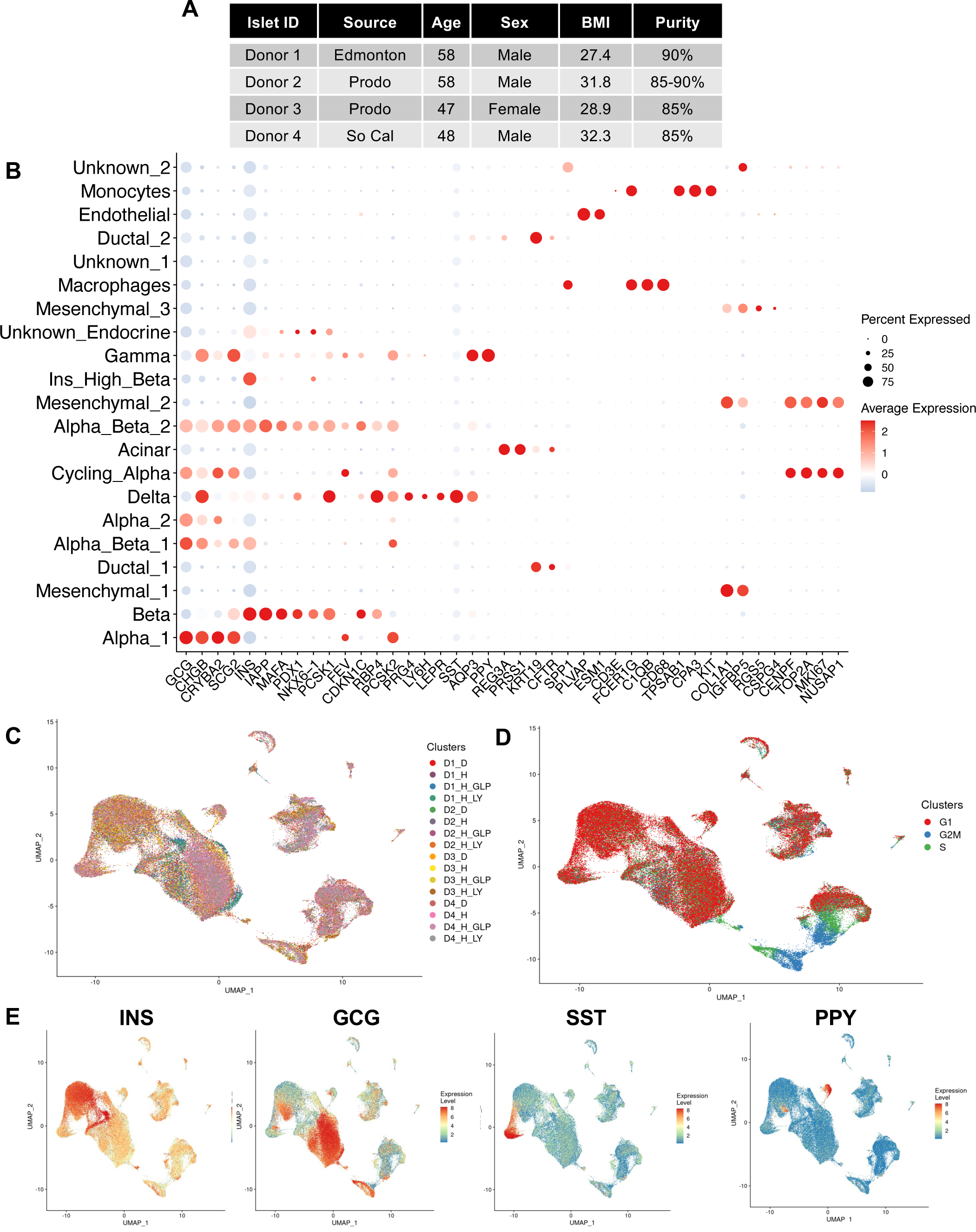
Presence of All Relevant Islet Cell Types In All Four Donors. **A.** Demographic information on the donors included in the study. **B** Dot plot showing normalized expression level and percent expressing cells for selected marker genes in each cluster. **C** UMAP plot showing cells labeled based on dataset name and **D** cell cycle phase. **E** Feature plots showing cells labeled with expression level of islet cell type hormones insulin (*INS*), glucagon (*GCG*), somatostatin (*SST*) and pancreatic polypeptide (*PPY*).

**Supplementary Figure 2.**
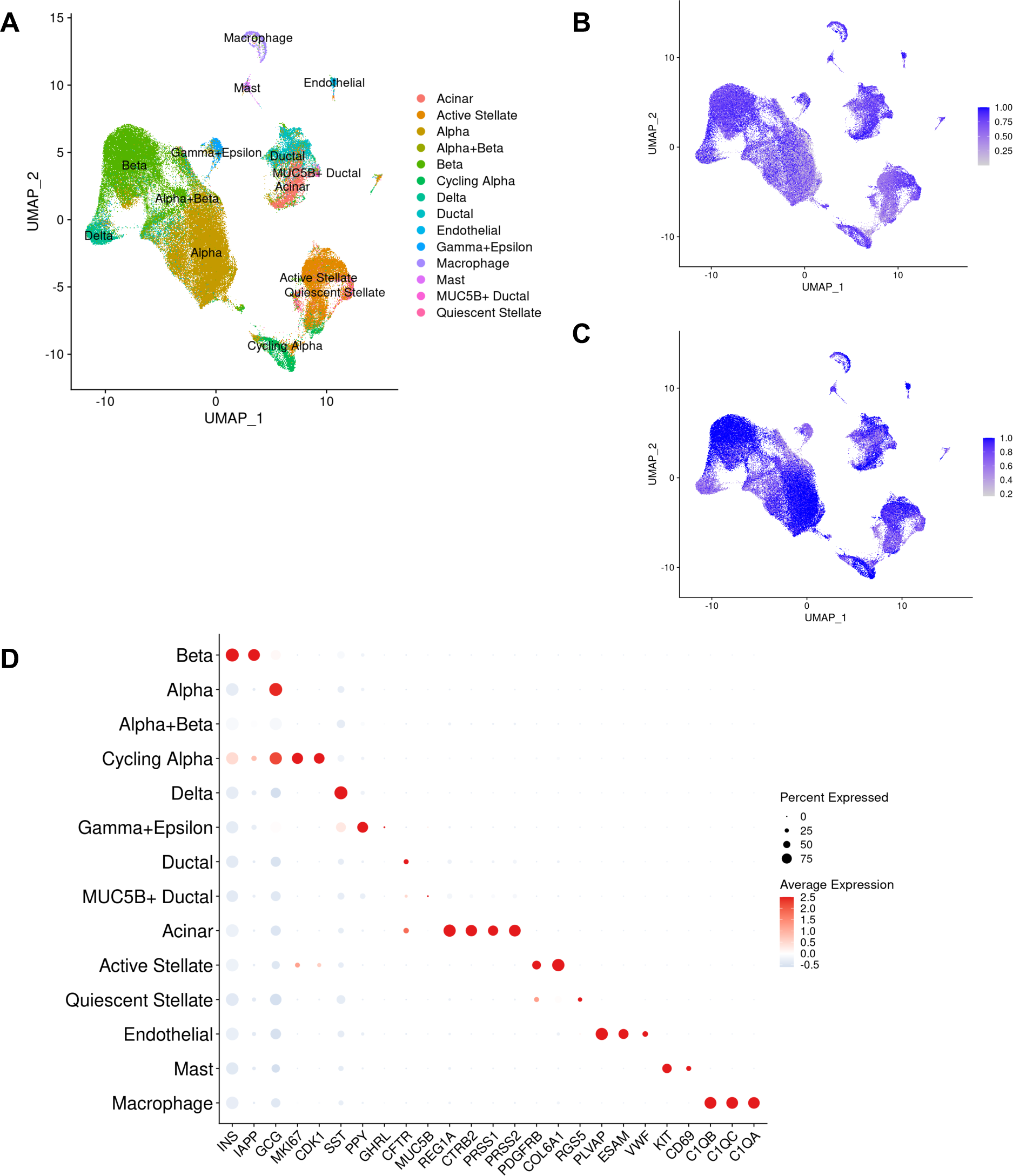
Concurrence of the Dataset Annotations with the Conventional Azimuth Database Annotations. **A** UMAP plot showing predcited cell annotation from HPAP human pancreas reference dataset. **B** UMAP plot showing mapping score on label transfer. **C** UMAP plot depicting predicted cell type score on label transfer. **D** Dot plot showing normalized average expression level and percent expressing cells for selected marker genes in each cluster from **A**.

**Supplementary Figure 3.**
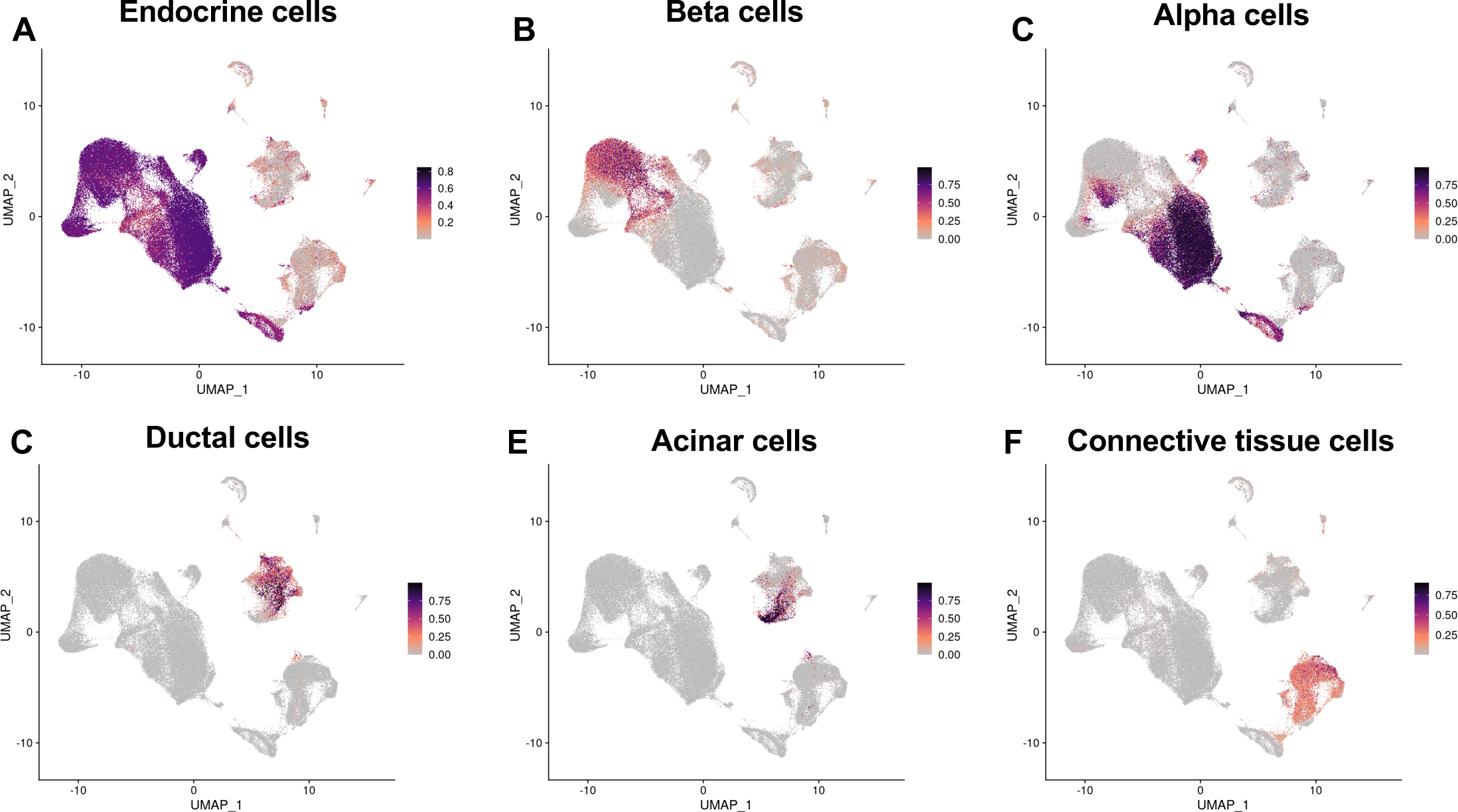
Corroboration of Annotations of Islet Cell Types With Unicell Database. Feature plots showing probability of **A** endocrine cells, **B** beta cells, **C** alpha cells, **D** ductal cells, **E** acinar cells, and **F** connective tissue cells based on Unicell database.

**Supplementary Figure 4.**
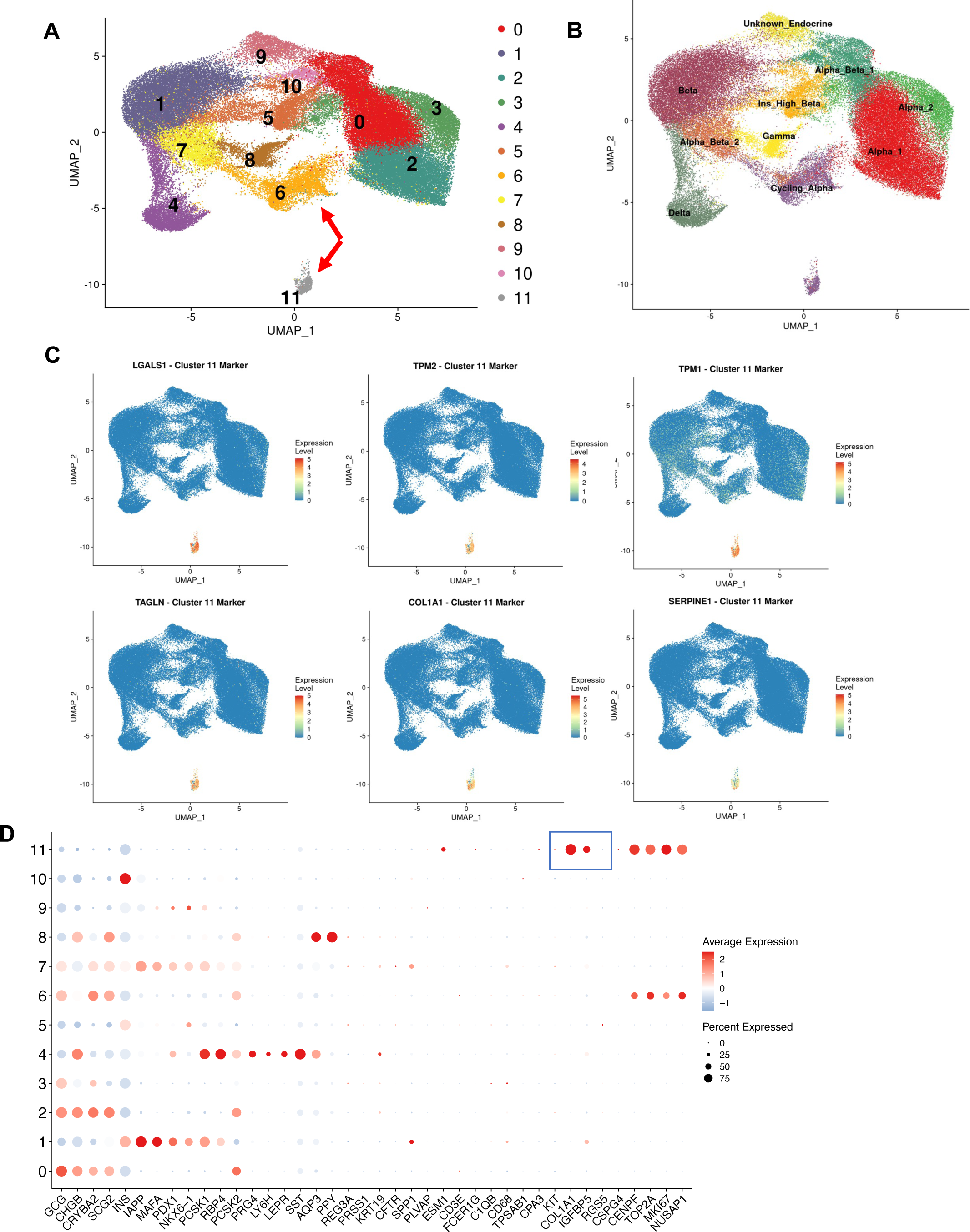
Cycling Alpha Cell Cluster Is Comprised of Endocrine and Non-Endocrine Cells. **A** UMAP plots showing post cell-cycle regression unsupervised clustering of single cell RNA-seq profiles of only endocrine cells. **B** Same UMAP plot as in A, depicting cell annotation projections. Red arrows point to two sub-clusters in the Cycling Alpha cluster. **C** Feature plots showing the expression of various genes in endocrine cells. Note that all genes shown here are highly expressed in sub-cluster 11. **D** Dot plot showing normalized expression level and percent expressing cells for selected marker genes in each cluster.

**Supplementary Figure 5.**
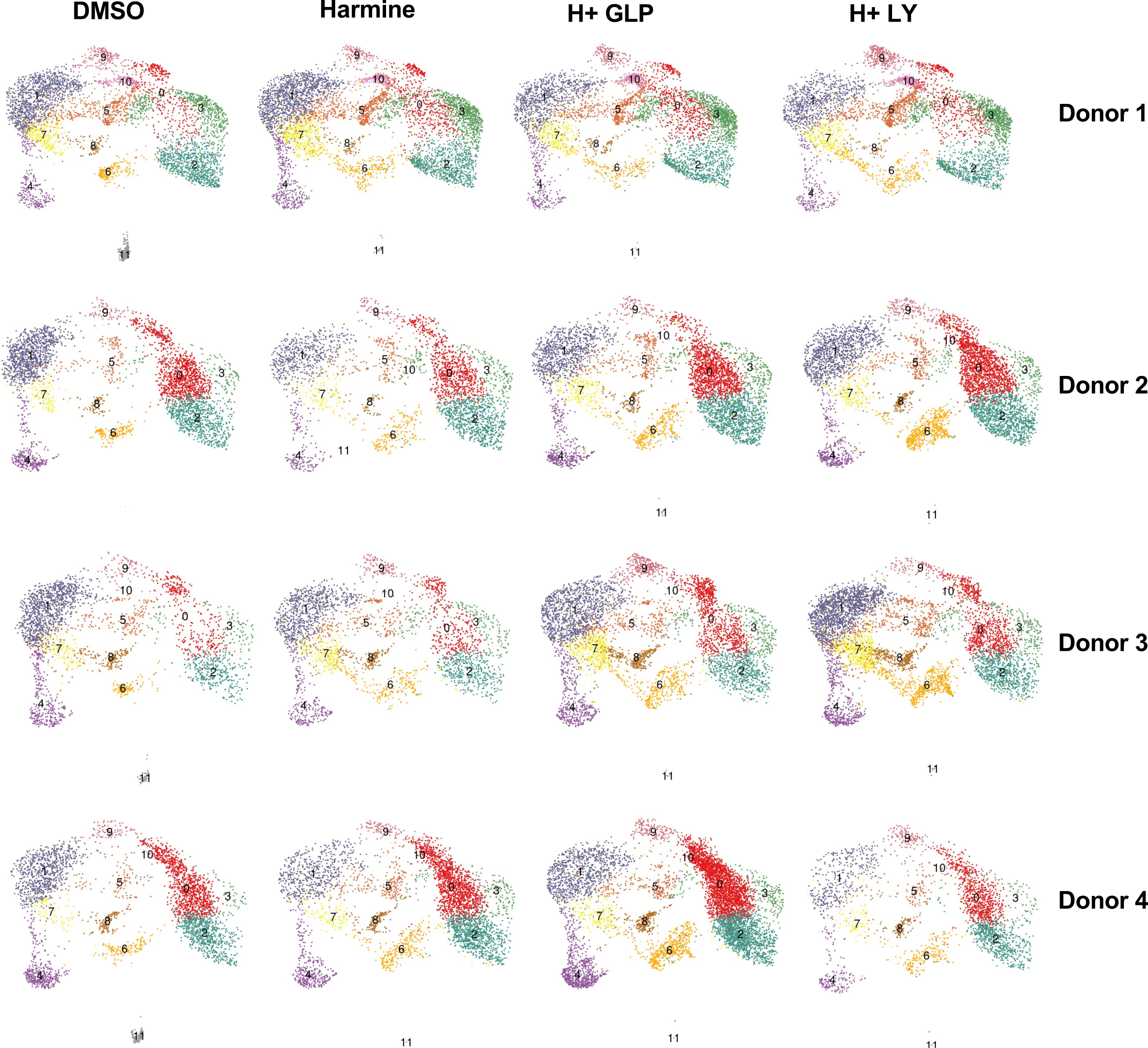
Analysis of Individual Human Islet Donors Confirms the Presence of Mesenchymal-Like Cells as Part of the Cycling Alpha Cluster. Split UMAP plots showing single cell RNA-seq profiles from each donor in all four treatment groups.

**Supplementary Figure 6.**
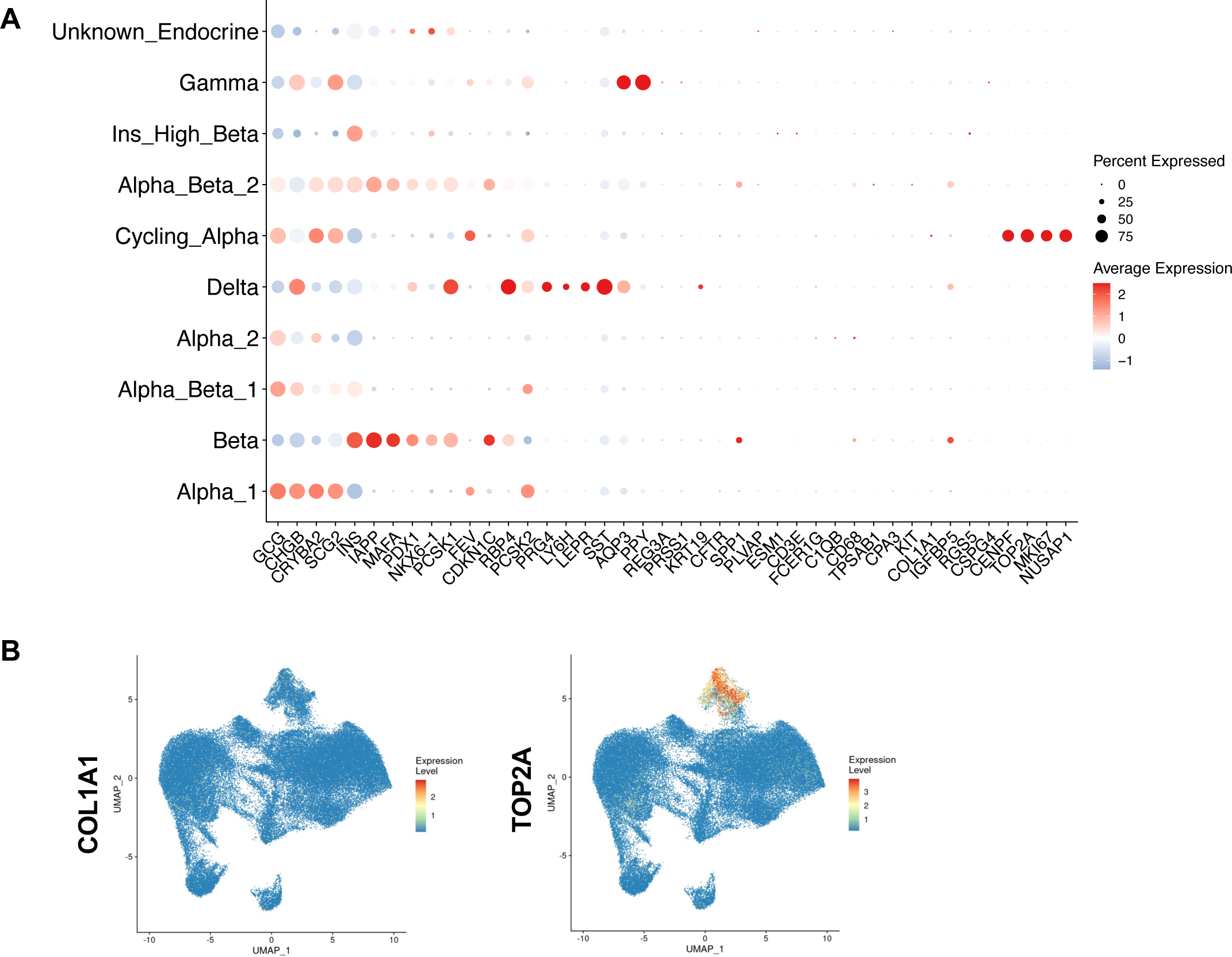
Confirmation of Removal of Non-Endocrine Cells from the Cycling Alpha Cell Cluster. **A** Dot plot showing normalized expression level and percent expressing cells for selected marker genes in each cluster. **B** Feature plots showing the expression of COL1A1 and TOP2A as a mesenchymal cell marker and a cell cycle marker, respectively.

**Supplementary Figure 7.**
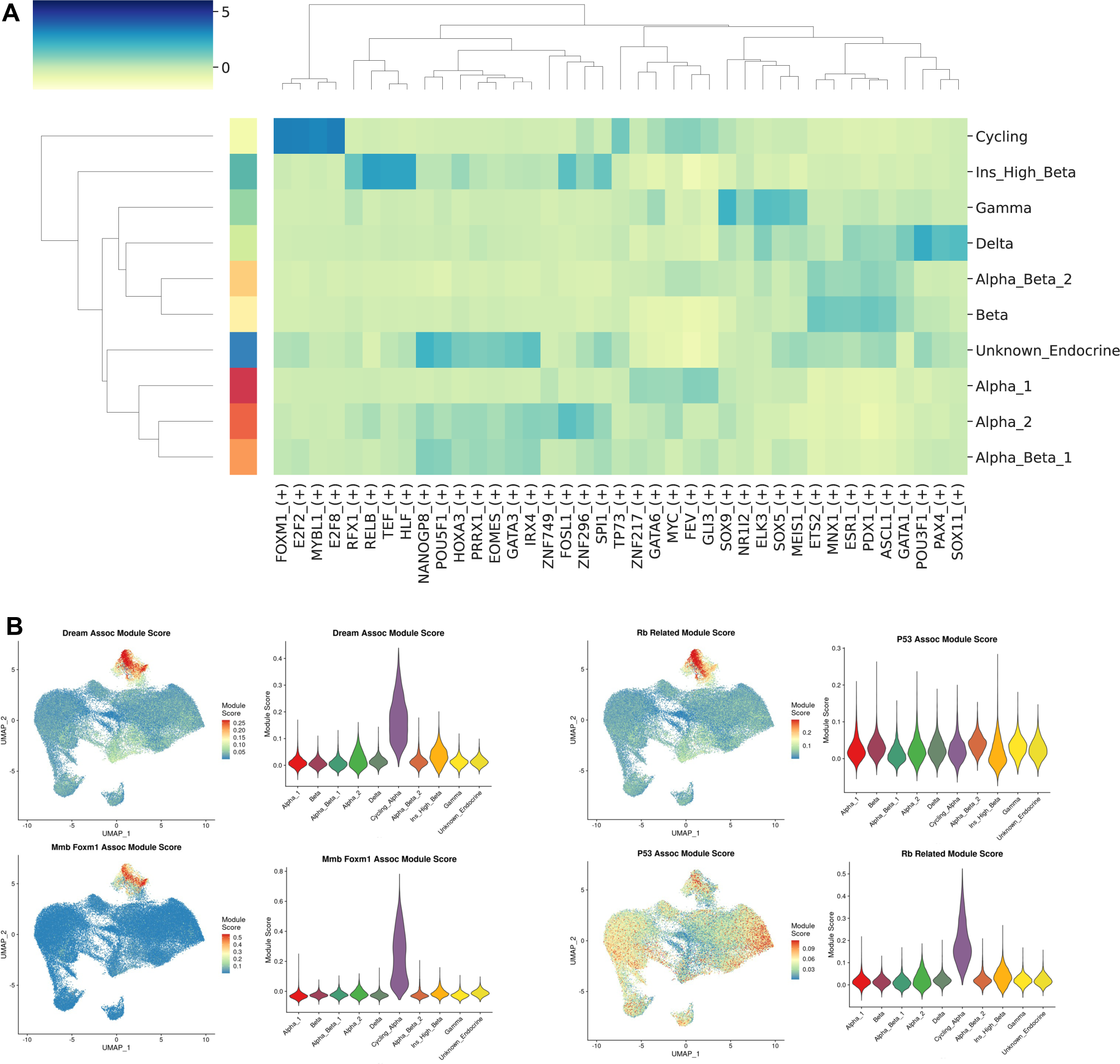
DREAM complex-associated transcriptional regulators are enriched in Cycling Alpha Cells. **A** Heatmap depicting the specificity score of top five highly enriched transcriptional regulators for each endocrine cell type. **B** Feature and bar plots demonstrate the enrichment level of Dream-, MMB-, RB- and P53-associated modules across endocrine cell types.

**Supplementary Figure 8.**
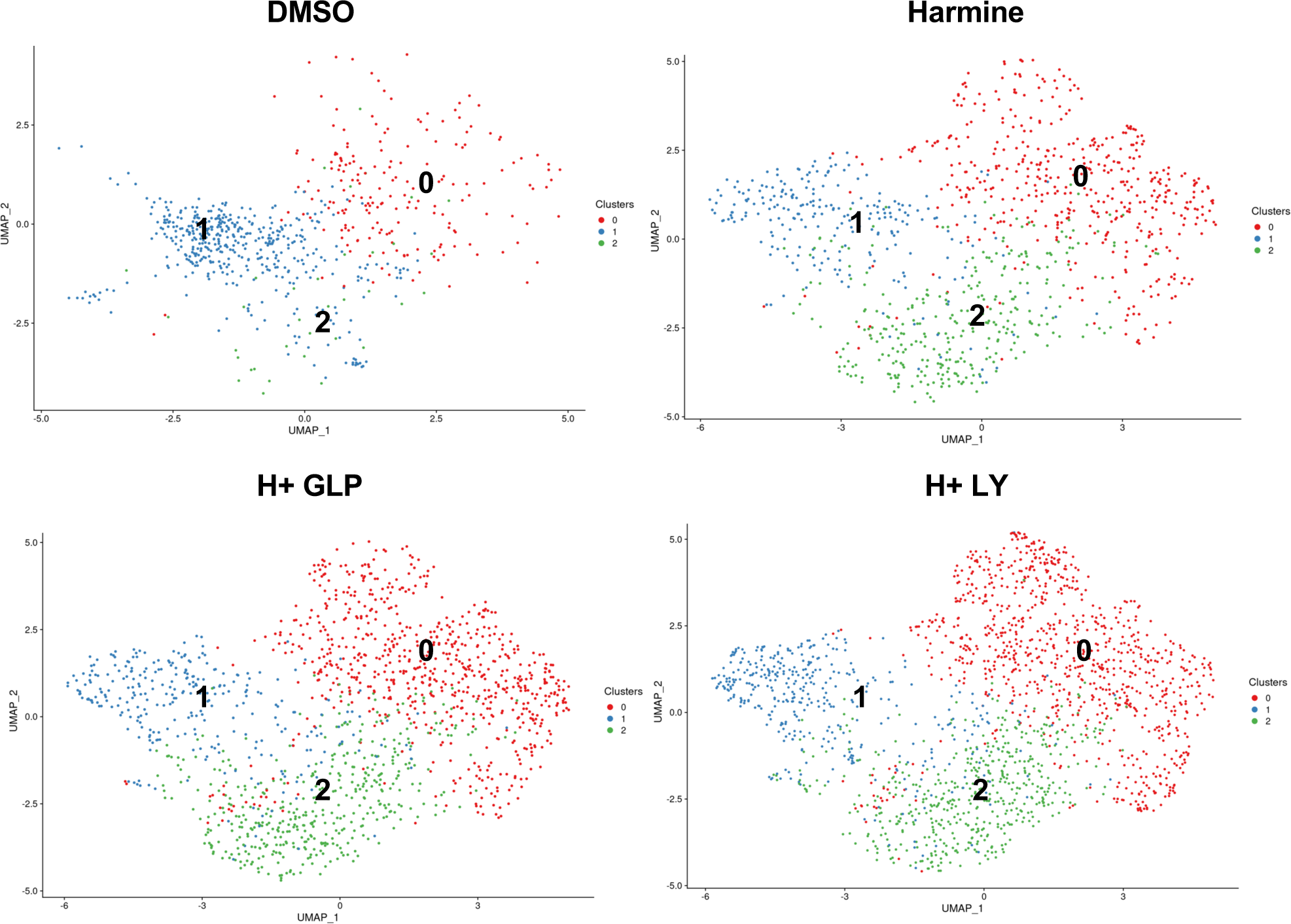
Analysis on the Sub-Clusters of Cycling Alpha Cells Reveal an Increase of Cycling Cell Proportion upon Regenerative Drug Treatment. Split UMAP plots showing single cell RNA-seq profiles from each treatment group.

